# Advancing Knotted Protein Design with ESM3: Guided Generation and Topological Insights

**DOI:** 10.64898/2026.05.07.723606

**Authors:** Eva Marsalkova, Petr Simecek

**Affiliations:** National Centre for Biomolecular Research & CEITEC Masaryk University, Brno, Czech Republic; Central European Institute of Technology (CEITEC) Masaryk University, Brno, Czech Republic

## Abstract

Multimodal protein language models have transformed protein design, yet their capacity to capture complex topological features remains poorly understood. We use knotted proteins, rare structures in which the backbone forms a nontrivial topological knot, as a test case to probe this capacity using ESM3, a generative protein language model. Topology-aware guided decoding strongly enriches ESM3 outputs for knotted topologies, producing structures classified as knotted at an 89% success rate (95% CI: 81– 94%), compared to ~0.5% for unguided diffusion-based approaches. A confidence analysis shows that freshly generated artificial knots have lower ESM3 pLDDT and pTM than real knotted proteins evaluated under the same pipeline, motivating a cautious interpretation of generated examples as model samples pending independent validation. In contrast, the robustness analyses on real knotted proteins are high-confidence: on average 84% of the protein sequence must be altered before the knot breaks, and the loss follows a sharp threshold rather than gradual degradation. Strikingly, structural drift accumulates well before topological disruption, suggesting that topology is more robust than specific three-dimensional arrangement. These findings position knotted proteins as a useful probe of how generative protein models represent rare, global structural features.

## 1 Introduction

Recent years have seen remarkable progress in AI-driven protein design. Structure prediction models such as AlphaFold2 [1] and ESMFold [11] predict three-dimensional structures with near-experimental accuracy, while generative tools including RFdiffusion [21], ProteinMPNN [6], and EvoDiff [2] enable the de novo design of proteins with desired structural properties. Multimodal foundation models further expand this design space: ESM3 jointly reasons over protein sequence, structure, and function [8]; ProTrek unifies protein sequence, structure, and natural-language function through contrastive learning [16]; and ProtST augments protein sequences with biomedical text descriptions of protein function and properties through multimodal representation alignment and multimodal mask prediction [22].

Despite these advances, an important question remains: to what extent do these models capture *topological* features of protein structure? Topology describes aspects of protein architecture that are more fundamental than specific atomic coordinates, namely properties preserved under continuous deformation. A particularly fascinating topological feature in proteins is the knot: a configuration in which the polypeptide backbone forms a nontrivial topological knot that cannot be untied without breaking covalent bonds [19, 20]. Knotted proteins are rare, occurring in less than 1% of known structures, yet they have been associated with enhanced kinetic stability [12, 18], resistance to proteolytic degradation [15], and distinctive functional roles. For example, the trefoil-knotted bacterial methyltransferase TrmD uses its knotted fold to couple AdoMet binding to tRNA binding and active-site assembly, thereby facilitating methyl transfer [3]. In ubiquitin C-terminal hydrolases, the deeply embedded 5_2_ knot has been proposed to protect the enzyme from unfolding and degradation by the proteasome [20], and mechanical unfolding experiments support the idea that knotted topology can impede processive degradation [15]. In transcarbamylases, comparative analyses suggest that the appearance of a knot coincided with a shift in enzymatic function [20]. Together with the strong evolutionary conservation of many knotted and slipknotted patterns, including conserved slipknots that may strap together helices in some transmembrane channels [17], these examples suggest that protein knots can contribute to biologically relevant structural and functional properties. Databases such as KnotProt [5] and AlphaKnot [13] catalogue the growing repertoire of known knotted topologies, while computational tools like Topoly [4] enable their detection using polynomial invariants. Experimentally validated de novo knotted proteins have also been reported, including crystallographic validation of a designed trefoil-knotted tandem-repeat protein [7]. Recent work by Klimentová and Simecek [9] demonstrated that diffusion-based protein design can generate artificial knotted proteins, though with a success rate of approximately 0.5%.

The ability of generative models to reconstruct complex protein structures from partial information also raises questions for biosecurity. It is common practice to redact or delete portions of sensitive protein sequences before publication to reduce dual-use risks [10]. Knotted proteins offer an ideal model system to study the effectiveness of such redaction: they are structurally complex, require precise spatial arrangements, have clear detection criteria, yet are not themselves harmful, allowing rigorous experimentation without biosafety concerns.

In this work, we leverage ESM3 to systematically investigate how protein language models interact with topological complexity. While we use only the smallest open-source variant, ESM3-SM with 1.4B parameters, we believe the results illustrate general principles about the relationship between learned protein representations and topology. We find that topology-aware guided decoding can strongly enrich ESM3 outputs for knotted topologies, producing diverse knot types from fully masked inputs, while generated sequences show no close similarity to known knotted proteins. Confidence analysis further indicates that generated examples require cautious interpretation and independent validation before being considered physical designs. We also observe that knot topology in real knotted proteins is remarkably robust to sequence perturbation, often persisting after extensive masking and regeneration even when the predicted three-dimensional structure has drifted substantially. These findings suggest that knotted proteins provide a useful test case for probing how generative protein models represent rare, global structural features.

## 2 Methods

### 2.1 Dataset

We use the real-protein subset of the publicly available `EvaKlimentova/Diffusion-all_knots` dataset [9]. This subset contains 5,000 proteins, comprising 1,000 knotted and 4,000 unknotted examples. The knotted and unknotted subsets have comparable sequence-length distributions: knotted proteins have a mean length of 399 residues (median 340, standard deviation 196, range 80–1000), while unknotted proteins have a mean length of 405 residues (median 339, standard deviation 204). This similarity reduces the possibility that downstream comparisons are driven primarily by protein length. For the knotted subset, knot types and knot-core positions were verified against AlphaKnot 2.0 [13]; the predominant topology is the trefoil knot (3_1_). The knot-core annotations were used in the targeted masking analyses described below.

### 2.2 ESM3 Model and Guided Generation

We employ ESM3-SM (1.4B parameters) [8] in half precision (fp16) on NVIDIA A10G GPUs via Modal serverless compute. ESM3 is a generative masked language model that reasons jointly across sequence, structure, and function tracks, enabling iterative unmasking of partially specified proteins. Given a partially masked protein, the model predicts probability distributions over tokens at masked positions; iterating this process progressively fills in the complete protein.

For guided generation, we use derivative-free guided decoding [14] as implemented in the ESM3 SDK. At each decoding step, multiple candidate unmaskings are sampled and scored by a user-defined objective; the highest-scoring candidate is kept, and the process repeats. We define a knot scoring function *s*(*x*) using the Topoly Python package [4] with Alexander polynomials. Topoly uses stochastic closure (projecting the open protein chain along random directions to form closed curves), yielding a probability distribution over topological types. With *N*_proj_ = 30 random projections, the score is:

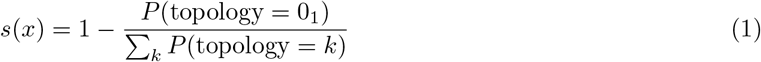

where 0_1_ denotes the trivial (unknotted) topology, and *P* (topology = *k*) is the fraction of stochastic closures yielding topology *k*. For de novo generation, we start from a fully masked sequence of length 256 and apply 8 decoding steps with 10 candidate samples per step, selecting the candidate with the highest knot score at each step.

### 2.3 Continuous Knot Score and Masking Strategies

We propose a continuous knot score based on randomized masking (Algorithm 1; the overall workflow is summarized in Supplementary Figure S1). For each protein, we randomly replace *X*% of residues with mask tokens, use ESM3 to regenerate the masked positions (8 generation steps, temperature 1.0), predict the structure of the resulting sequence (8 structure generation steps), and evaluate topology via Topoly. Averaging over *N* = 8 independent trials yields a score in [0, 1]. We systematically vary *X* from 10% to 90%.

We compare three masking strategies: **random masking** (uniformly sampled positions), **contiguous masking** (a single block), and **targeted masking** of the knot core versus flanking regions, using ground-truth core positions from AlphaKnot 2.0.

#### Algorithm 1

Randomized Knot Score Calculation

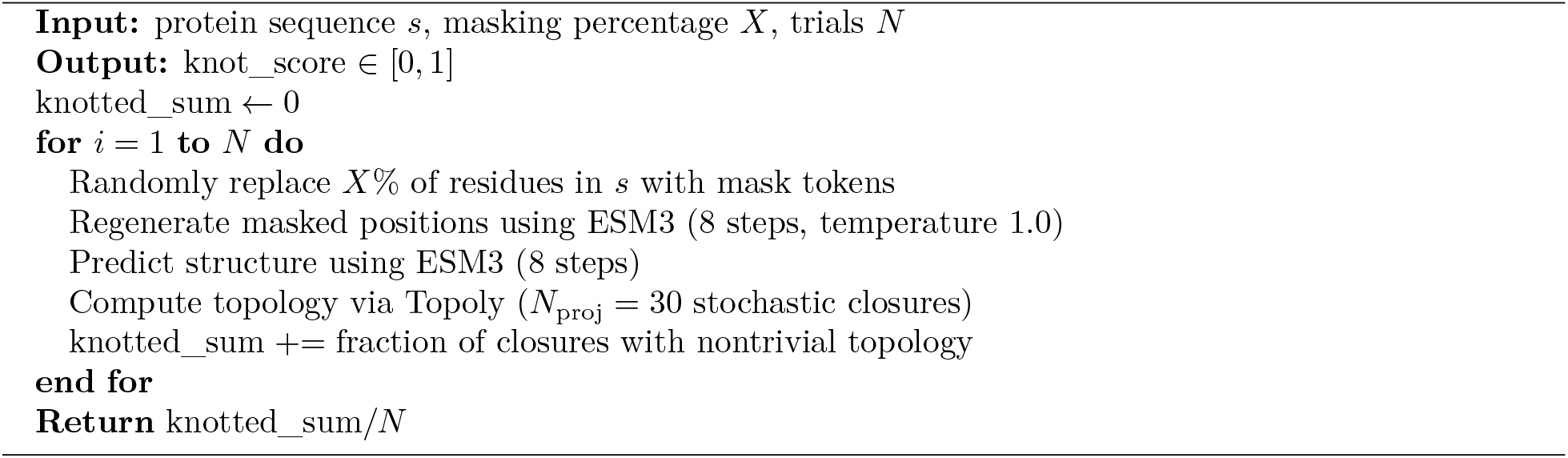

### 2.4 Structural Drift and Embedding Analysis

We measure backbone RMSD between the ESM3-predicted structure of the original sequence and the ESM3-predicted structure after masking and regeneration, using the Aligner utility from the ESM3 package (C*α* atom superposition). The RMSD analysis uses a smaller sample (*n* = 80) than the masking analysis (*n* = 250) because each RMSD measurement requires generating and aligning two full structures, making it computationally more expensive. We also compute pairwise sequence identity as the fraction of matching residues at aligned positions, normalized by the length of the longer sequence.

For embedding analysis, we extract mean-pooled ESM3 embeddings (dimension 1,536) from protein sequences and train a multilayer perceptron with two hidden layers (1,024 and 256 neurons, ReLU activation) for binary classification (knotted vs. unknotted), using an 80/20 stratified train/validation split (4,000 train, 1,000 validation) with early stopping on validation loss (scikit-learn MLPClassifier).

## 3 Results

### 3.1 Guided Generation of Knotted Proteins

ESM3’s guided generation achieved an 89% success rate (89/100; 95% CI: 81–94%) in producing outputs classified as knotted from fully masked sequences, compared to the ~0.5% success rate reported for unguided diffusion-based approaches [9]. We note that this comparison is between guided and unguided generation and should be interpreted accordingly: the improvement reflects the effectiveness of guided decoding with a topology-aware scoring function within the ESM3–Topoly evaluation pipeline, not a direct comparison of model architectures or a validation of physical foldability.

Among the 70 proteins with resolved knot types (the remaining 19 exceeded the Alexander polynomial’s default crossing limit of 15 and could not be classified; see Limitations), trefoil knots (3_1_) were most common (33 occurrences), followed by figure-eight knots (4_1_, 10) and torus knots (5_2_, 10). More complex types including 8_19_, 9_44_, and 9_45_ were also generated (Figure 2).

**Figure 1:**
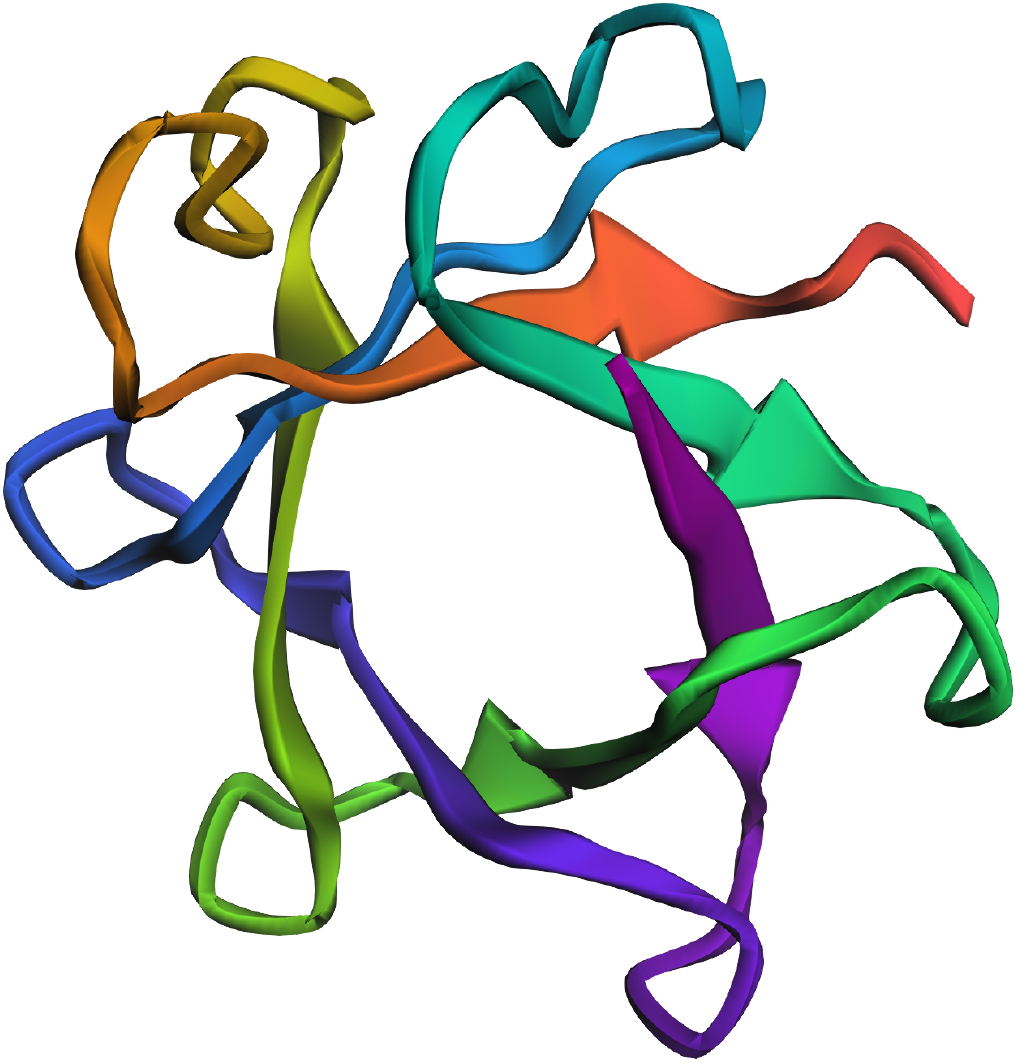
Rainbow-colored ribbon-cartoon rendering of the AlphaKnot model for the knotted protein A0A165M1R5 (*Acidovorax* sp.), featuring a 7_1_ knot. The backbone is colored from N-terminus to C-terminus to make the chain path easier to follow. Interactive 3D visualization: https://alphaknot.cent.uw.edu.pl/view/A0A165M1R5/1/E1.

**Figure 2:**
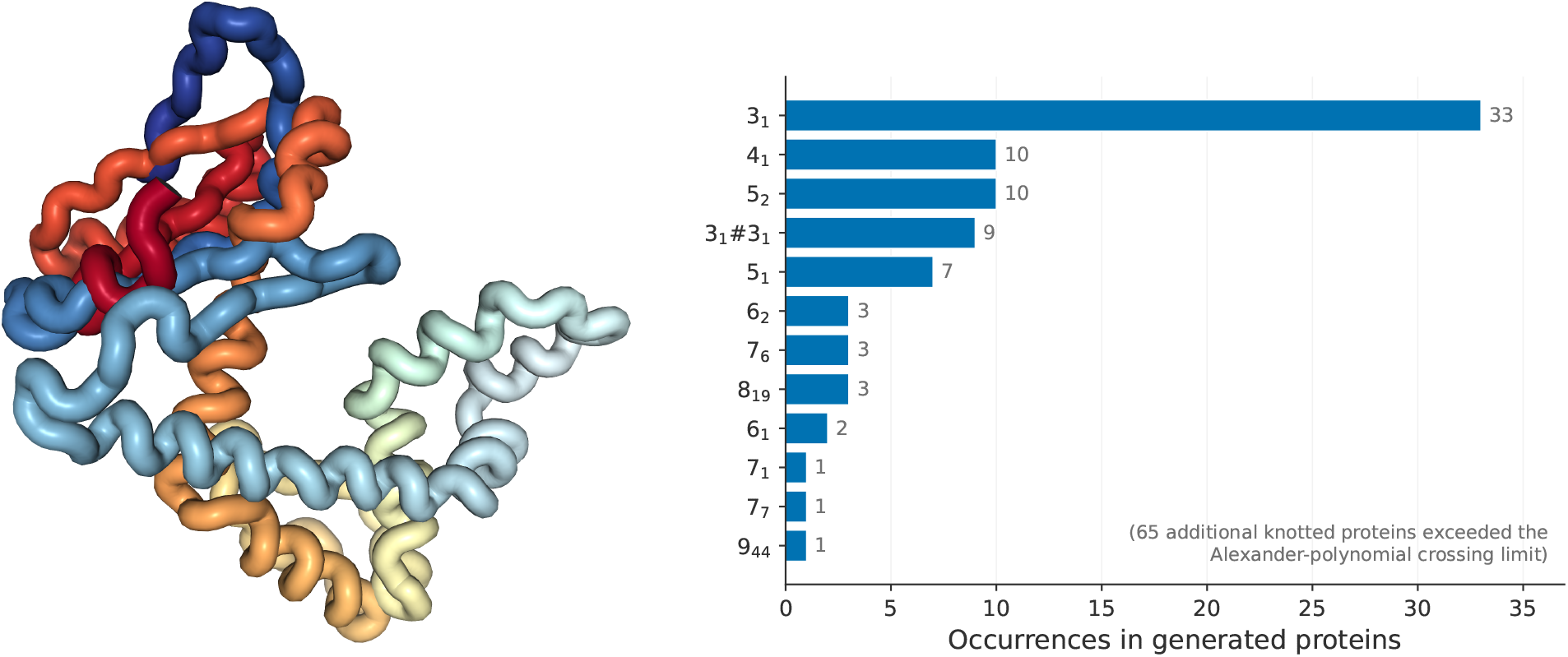
Left: rainbow tube rendering of an example topology-guided ESM3 output classified as knotted, colored by residue index along the chain to make the backbone path easier to follow (knot score 1.0; the rendered ESM3-predicted structure has mean pLDDT 0.51 and pTM 0.33). The moderate confidence reinforces that generated examples should be interpreted as topology-enriched model samples rather than validated folds. Right: distribution of resolved knot types in successfully generated proteins. The trefoil (3_1_), the simplest and most common knot in natural proteins, dominates, but complex topologies with up to 9 crossings are also produced.

We further tested knot-type-specific generation by modifying the scoring function to preferentially reward a particular topology. Targeting the trefoil was highly effective (9/10 hit the target), consistent with 3_1_ being the simplest and most common knot type. When the model generates a knot but misses the target, it naturally defaults to the most probable topology. More complex targets (4_1_, 5_1_, 5_2_) were achieved in 20–30% of attempts, with generation biased toward trefoils in the remaining cases (Supplementary Figure S10). The success rate also depends on target protein length: at 100 residues only 20% of attempts produced knots, rising to 90% at 200 residues and 100% at 350 or more (Figure 3), consistent with the observation that natural knotted proteins are typically 200–400 residues long.

**Figure 3:**
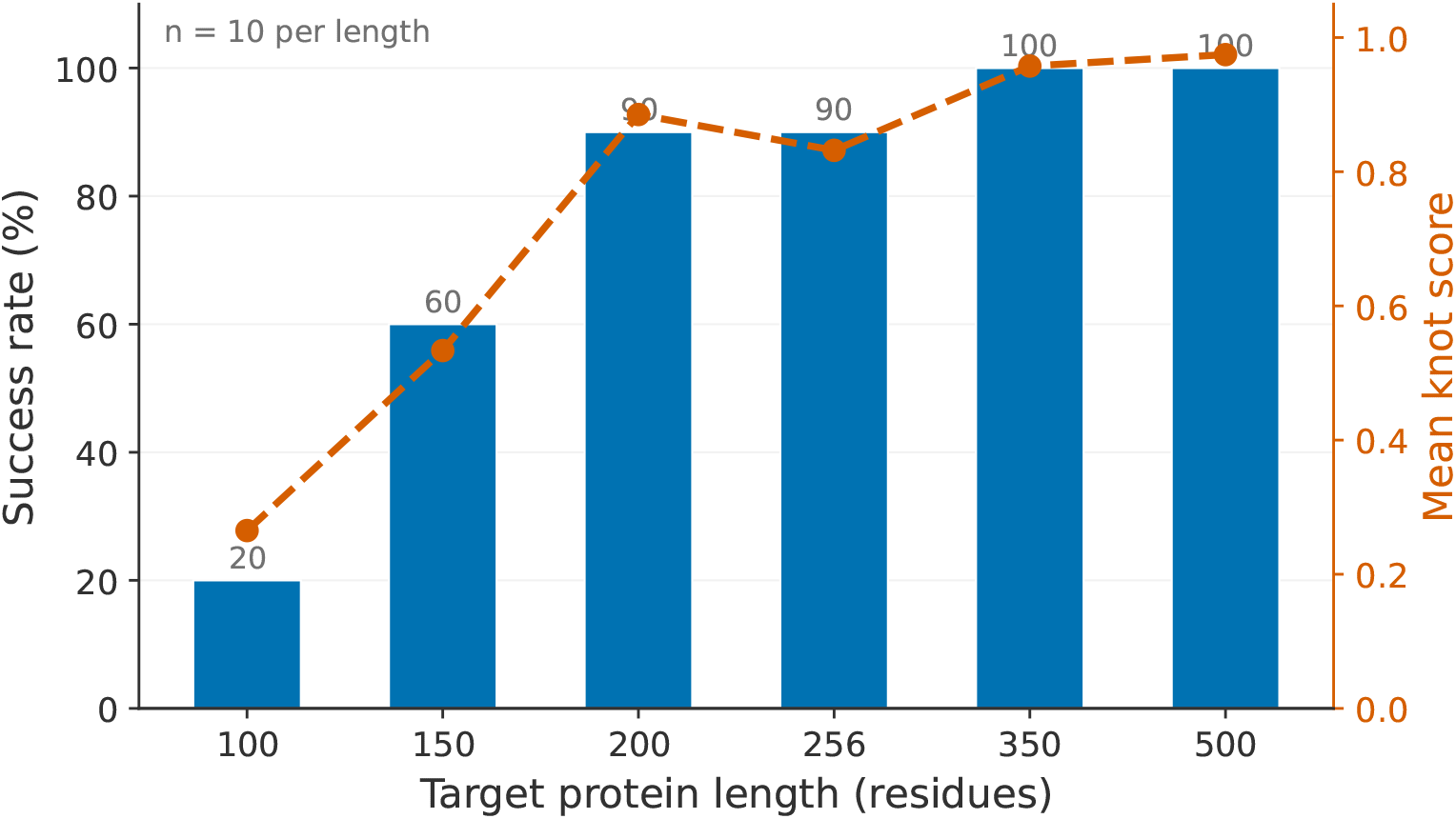
Guided generation success rate as a function of target protein length (*n* = 10 per length). Bars (left axis) show the fraction of attempts that produced a knot; the orange dashed line (right axis) shows the mean knot score. Generation is unreliable below 150 residues but saturates at near 100% from 200 residues onward, consistent with the typical length range of natural knotted proteins.

### 3.2 Confidence of ESM3-Predicted Structures

Because topology alone does not establish physical plausibility, we performed an additional confidence analysis using ESM3’s own structure-confidence outputs. In a fresh guided-generation replicate (*n* = 20), 16/20 generated proteins were classified as knotted by Topoly (mean knot score 0.807), with lower confidence scores than observed for real knotted proteins (mean pLDDT 0.317, median 0.281; mean pTM 0.210, median 0.162). As a baseline, we applied the same ESM3 structure-prediction and topology-evaluation pipeline to 20 real knotted proteins from the dataset; these retained high knot scores (mean 0.892) and had high confidence (mean pLDDT 0.895, median 0.933; mean pTM 0.841, median 0.898; Supplementary Figure S2).

These results indicate that topology-aware guided decoding strongly enriches for structures that ESM3 and Topoly classify as knotted, while also showing that generated examples should be treated as model-derived candidates pending independent validation. We therefore interpret the guided-generation results primarily as evidence that ESM3 can be steered toward rare topological states in its learned sequence– structure space.

A simple MLP classifier trained on mean-pooled ESM3 embeddings achieves 97.1% accuracy in distinguishing knotted from unknotted proteins (*n*_val_ = 1000; Supplementary Figure S7), suggesting that ESM3’s learned representations encode information relevant to topology even without explicit topological supervision.

### 3.3 Knot Robustness and Sharp Threshold Behavior

Our masking stability analysis (*n* = 250 proteins, 10 masking levels, 8 trials each) reveals that knot topology is remarkably stable under sequence perturbation (Figure 4). At 50% masking, 94% of knotted proteins retain their topology. At 70% masking, where only 30% of the original sequence remains, 85% are still knotted. The mean breaking point is 84% (± 1.2% SE).

**Figure 4:**
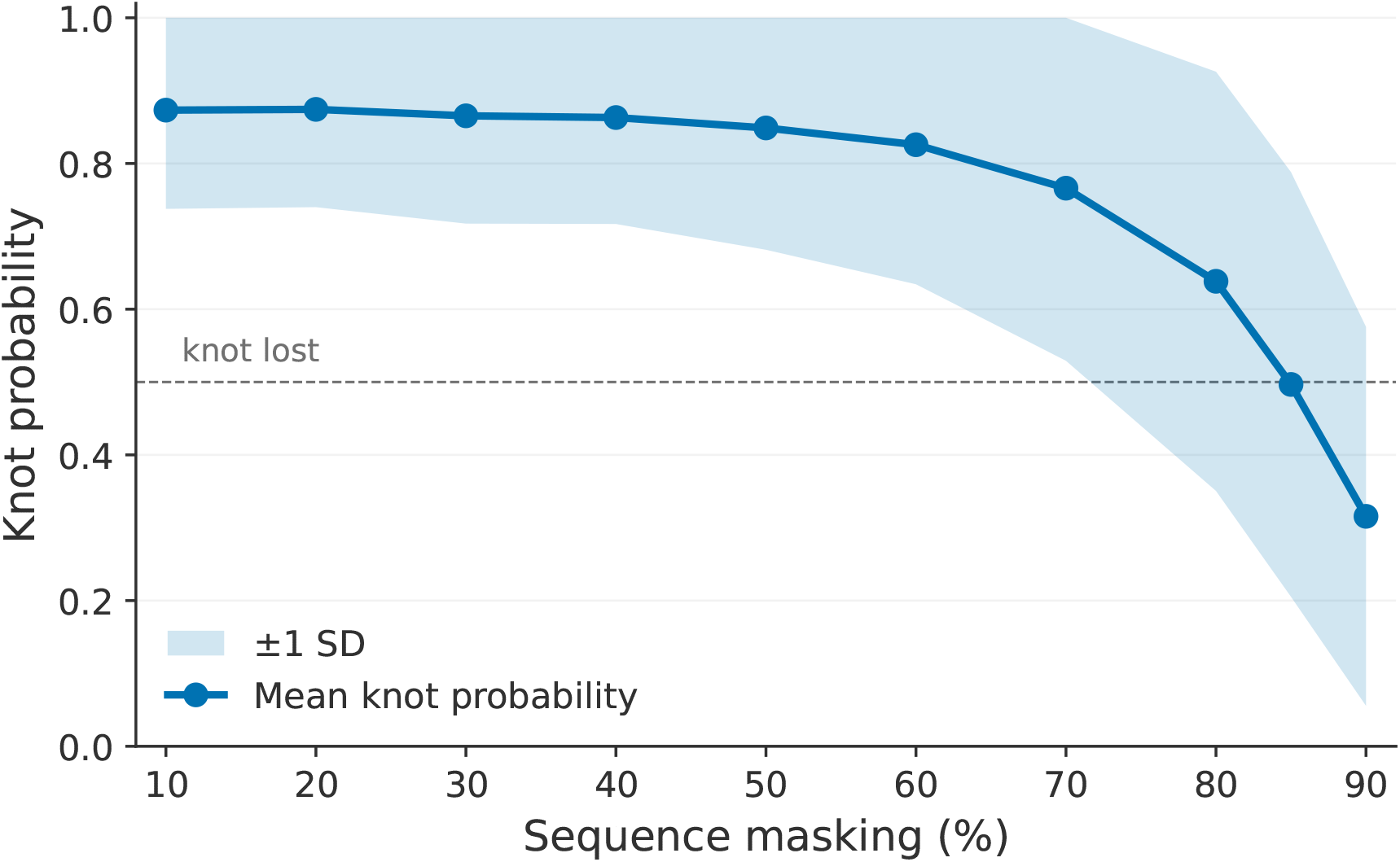
Knot probability as a function of masking percentage (*n* = 250). Shading shows ± 1 standard deviation. The knot persists until approximately 80–90% of the sequence is masked, then drops sharply.

The loss of topology appears abrupt rather than gradual: 68% of proteins exhibit a single-step drop in knot probability exceeding 0.3 between consecutive masking levels (Supplementary Figure S3). While the 10-percentage-point spacing of our masking grid limits our ability to distinguish a true discontinuity from a steep sigmoid, the behavior is consistent with a sharp threshold-like transition, suggesting a critical amount of sequence information is required to specify the topology.

### 3.4 Structural Drift Precedes Topological Disruption

The RMSD analysis (*n* = 80; smaller than the masking analysis due to the additional computational cost of structure generation and alignment) reveals a key decoupling between structural conservation and topological persistence (Figure 5). At 50% masking, the median RMSD is already 3.24 Å and sequence identity has dropped to 73%, yet the mean knot probability remains 0.85. At 70% masking, RMSD reaches 4.58 Å (substantial structural rearrangement), while the knot probability is still 0.76.

**Figure 5:**
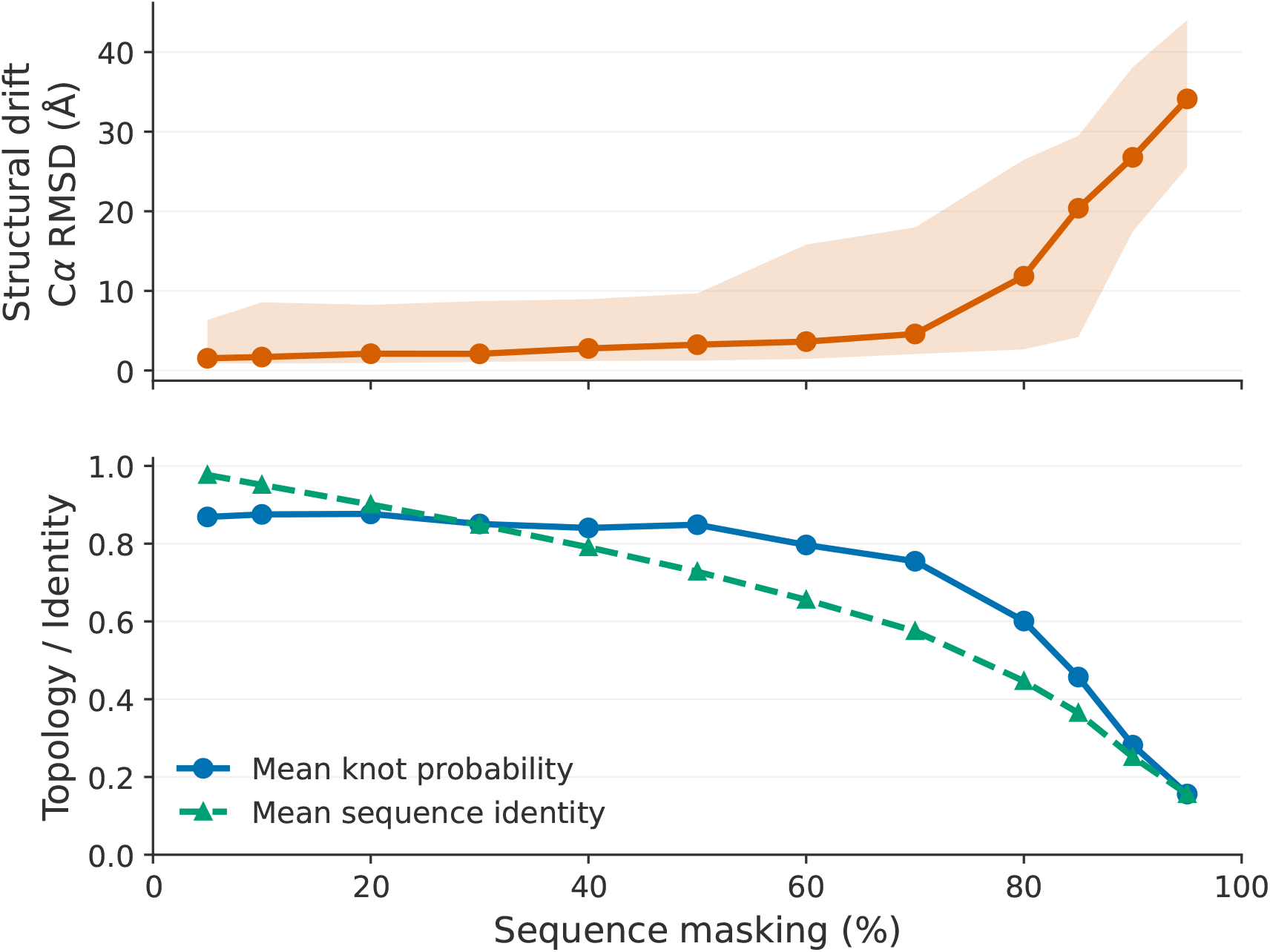
Top: structural drift (median C*α* RMSD) accumulates steadily with masking, with shading showing the 25–75% inter-quartile range across proteins. Bottom: knot probability (solid) persists until much higher masking levels, while sequence identity (dashed) declines monotonically. The decoupling between the two panels suggests that topology is more robust than specific 3D arrangement.

This decoupling suggests that topology is more robust than specific three-dimensional arrangement. The model generates structures that differ substantially from the original in atomic coordinates yet maintain the same topological class, consistent with the idea that the knot is encoded in a distributed manner across the sequence, with many different conformations capable of realizing the same topology.

### 3.5 Evidence Against Simple Memorization

One might worry that the model simply memorizes and retrieves knotted sequences from its training data rather than capturing nontrivial information about topology. Several lines of evidence argue against this interpretation.

First, de novo generation starts from a fully masked sequence, so there is no input information to trigger retrieval. Second, the generated knotted proteins show no close sequence similarity to known knotted proteins (Figure 6): the maximum identity between any generated protein and any of the 1,000 known knotted proteins in the dataset is just 14.5%, with a mean of 11.6%, indistinguishable from a random baseline of 12.5% (random unknotted proteins compared to the same knotted set). Not a single generated protein exceeds 15% identity with any known knotted protein. As an additional database-level check, we searched the 16 knotted outputs from the fresh confidence replicate against Swiss-Prot using BLASTP through the EMBL-EBI service; none returned a detectable hit under the service settings used. This supports sequence-level novelty, while the confidence analysis above cautions that novelty should not be conflated with physical plausibility.

**Figure 6:**
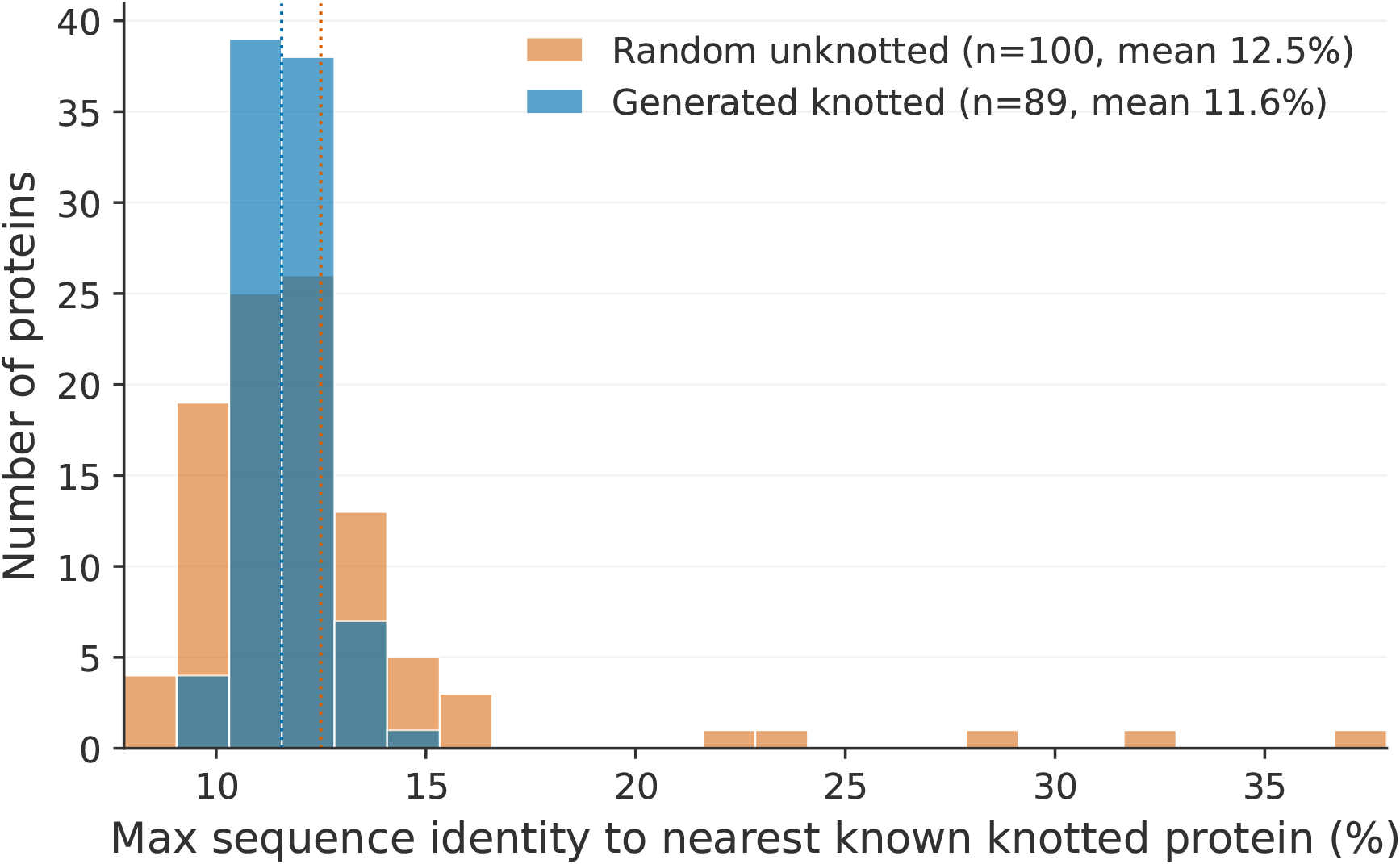
Maximum sequence identity of each generated knotted protein to its nearest known knotted protein (blue), compared to random unknotted proteins (orange baseline). Dotted vertical lines show the per-distribution means. The two distributions overlap, indicating that generated proteins are no more similar to known knotted proteins than random sequences are.

Third, at 50% masking the regenerated sequences share only 73% identity with the originals yet maintain 85% knot probability (Supplementary Figure S4). We note that low sequence identity does not by itself exclude structural similarity (two proteins can share very different sequences yet similar folds); however, the combination of novel sequence, novel generation context (all-mask input), and the ability to steer generation toward specific knot types via modified scoring functions collectively argues against simple template retrieval.

### 3.6 Unknotted-to-Knotted Conversion

Using iterative guided generation with low-rate masking (3% per iteration, up to 15 iterations), we converted 17 out of 99 unknotted proteins (≤250 residues) into knotted variants, achieving a 17% success rate (95% CI: 10–26%). This is substantially higher than the ~0.5% rate achievable without guided generation [9]. Successful conversions showed a bimodal pattern: 8 proteins converted within 3 iterations, while 9 required 7–14 iterations. The remaining 82 proteins showed almost no progress (61 scored exactly 0.0), suggesting that most unknotted proteins are strongly resistant to topological transformation. Beyond the confidence analysis reported above for de novo generation, we did not separately assess the physical plausibility of the converted structures, which remains an important direction for future work.

### 3.7 Supplementary Analyses of Masking and Representation Structure

The supplementary analyses provide additional context for the robustness results. Targeted masking of the annotated knot core reduced knot probability more strongly than masking non-core regions, but did not abolish topology: even when the full annotated core, corresponding to approximately 47% of the sequence, was masked, the mean knot probability remained 0.659 (Supplementary Figure S5; Supplementary Table S1). This indicates that the knot is not specified solely by the annotated core residues, but can often be reconstructed from broader sequence context. Consistent with this distributed encoding, the sliding-window analysis did not reveal a single uniquely vulnerable region along the sequence (Supplementary Figure S8).

The comparison of masking strategies further supports this interpretation. Contiguous masking was generally less disruptive than random masking at the same masking fraction (Supplementary Figure S6), suggesting that topology is more sensitive to dispersed loss of sequence information than to deletion of one local block. Individual transition curves and the breaking-point distribution show that most proteins retain high knot probability until very high masking levels, followed by a rapid decline near the breaking point (Supplementary Figures S3 and S9). Finally, embedding-based analyses show that knotted and unknotted proteins are readily separable in mean-pooled ESM3 representation space: the MLP classifier reaches 97.1% validation accuracy, and the UMAP projection shows partial but visible class structure (Supplementary Figure S7). Together, these supplementary analyses support the conclusion that topology-relevant information is distributed across sequence and reflected in ESM3 representations.

## 4 Discussion

Our results suggest that multimodal protein language models capture nontrivial information about protein topology, even in their smallest variants. The 89% success rate in topology-aware guided decoding, combined with the diversity of generated topologies, indicates that ESM3 can be effectively steered toward rare and complex topological states in its learned sequence–structure space. The confidence analysis shows that such steering is not by itself equivalent to validated protein design, but it provides a useful starting point for future confidence-aware or independently validated design workflows.

The sharp threshold behavior in knot stability is perhaps our most robust finding. Rather than a gradual degradation, most real knotted proteins show an abrupt loss of topology at a critical masking threshold. One interpretation is that topological information is distributed and partially redundant across the sequence: above a critical information threshold the model can reconstruct the knot; below it, the topology is rapidly lost. The decoupling between RMSD and topology, where structures diverge substantially in 3D space while maintaining the same knot type, further supports the view that topology is a more fundamental feature than specific atomic arrangement, consistent with the mathematical nature of topological invariants.

The confidence analysis also clarifies the circularity risk inherent in using the same model family for generation and structure evaluation. ESM3-SM assigns high confidence to real knotted proteins under our pipeline, while fresh guided-generation outputs have lower confidence on average. This internal check supports the use of ESM3 for probing topology-relevant representations, while motivating additional confidence-aware objectives and independent structure predictors in future design-oriented work.

These findings also have potential implications for biosecurity. Current practices of redacting portions of sensitive protein sequences assume that partial information is insufficient for reconstruction. Our results show that after masking 70% of a sequence, modern generative models can still recover complex topological features with 85% reliability. Targeted deletion of the knot core still allows reconstruction 66% of the time from flanking context alone (Supplementary Table S1). We emphasize several caveats: our experiments use random and block masking, which differs from real-world targeted redaction of specific functional regions; the threat model is not fully developed; and we do not claim that knotted proteins themselves pose biosecurity risks. Rather, the results suggest that the effectiveness of sequence redaction as a general safeguard deserves further scrutiny in the context of increasingly capable generative models.

### Limitations

Our experiments use ESM3-SM (1.4B parameters), the smallest publicly available variant; larger models (7B, 98B) would likely show stronger capabilities. All topology detection is computational (Alexander polynomials via stochastic closure), and we did not perform wet-lab validation of generated proteins. Although ESM3 assigns high confidence to real knotted proteins under the same structure-prediction pipeline, fresh guided-generation outputs had lower pLDDT and pTM on average. Thus, these generated structures should be interpreted as topology-enriched model samples pending independent structure prediction and experimental validation. Approximately 21% of generated knotted proteins (19/89) had topologies too complex for the Alexander polynomial to resolve within the default crossing limit of 15; using HOMFLYPT or Jones polynomials could potentially disambiguate these cases. The unknotted-to-knotted conversion rate (17%) is modest, and the bimodal success pattern suggests fundamental limitations of iterative guided generation for topological transformation.

**Table 1:** Summary of key results.

| Experiment | Result | 95% CI |
| --- | --- | --- |
| Guided generation success rate | 89% ($n = 100$) | [81, 94%] |
| Mean pLDDT, generated vs. real knots | 0.32 vs. 0.89 ($n = 20$ each) | — |
| Mean breaking point (masking) | 84% ($n = 250$) | $\pm 1.2\%$ SE |
| Knotted at 70% masking | 85% of proteins | — |
| RMSD at 50% masking | 3.24 Å median ($n = 80$) | — |
| Max seq. identity to known knotted | 14.5% ($n = 89$) | — |
| Embedding classifier accuracy | 97.1% ($n_{\text{val}} = 1000$) | — |
| Unknotted-to-knotted conversion | 17% ($n = 99$) | [10, 26%] |

### What this study does not show

We do not demonstrate that ESM3 “understands” topology in a mathematical sense. The model may have learned statistical regularities in sequence–structure relationships that happen to correlate with topological features, without representing topology explicitly. We also do not demonstrate biosecurity risks from knotted proteins specifically; they serve as a model system. Extension of these findings to actual dual-use scenarios would require additional analysis with realistic threat models.

### Future work

Key directions include experimental validation of generated knotted proteins, extension to other complex structural features (slipknots, lasso motifs [5]), systematic study of what sequence features make a protein amenable to topological transformation, and information-theoretic analysis of the minimum sequence information required to specify a given topology. Testing with larger ESM3 variants as they become publicly available would clarify whether the observed capabilities scale with model size.

## 5 Conclusion

We have investigated the interaction between multimodal protein language models and topological complexity, using knotted proteins as a test case. Our key findings are:

1. Topology-aware ESM3-guided decoding enriches for outputs classified as knotted with an 89% success rate and diverse topology, compared to ~0.5% for unguided approaches; a fresh confidence replicate shows that these generated examples require further validation before being interpreted as physical designs.
2. Knot topology in real knotted proteins exhibits a sharp threshold-like behavior under sequence perturbation: on average 84% of the sequence must be altered before the knot breaks.
3. Structural drift (RMSD) precedes topological disruption, suggesting that topology is more robust than specific 3D arrangement.
4. Generated knotted proteins show no close sequence similarity to known knotted proteins (mean 11.6%, indistinguishable from random), arguing against simple memorization.

These results are consistent with the view that protein language models capture nontrivial information related to topology. They also show why topology-aware generation must be paired with confidence assessment and, ultimately, independent validation before being interpreted as protein design. As such models continue to scale, understanding their topological capabilities becomes increasingly important, both for harnessing their potential in protein engineering and for anticipating challenges in biosafety governance.

## Supporting information

suppl. immages

## Data and Code Availability

All code used in this study is publicly available at https://github.com/ML-Bioinfo-CEITEC/KPDwESM3. The sequence-level dataset and associated metadata used in this study are available through the HuggingFace Hub at https://huggingface.co/datasets/EvaKlimentova/Diffusion-all_knots. The corresponding PDB structure files are archived separately on Zenodo at https://doi.org/10.5281/zenodo.21206269.

## Acknowledgments

The project was supported by the OPUS LAP program of the Czech Science Foundation, project no. 23-04260L (“Biological code of knots – identification of knotted patterns in biomolecules via AI approach”). Computational resources were provided by Modal serverless GPU compute, using $118.09 in free credits.

## Notes

https://github.com/ML-Bioinfo-CEITEC/KPDwESM3

### Competing Interest Statement

The authors have declared no competing interest.

### Summary of Updates

revision of images, better positioning of the paper

