## Supplementary material for "Advancing Knotted Protein Design with ESM3: Guided Generation and Topological Insights": suppl. immages

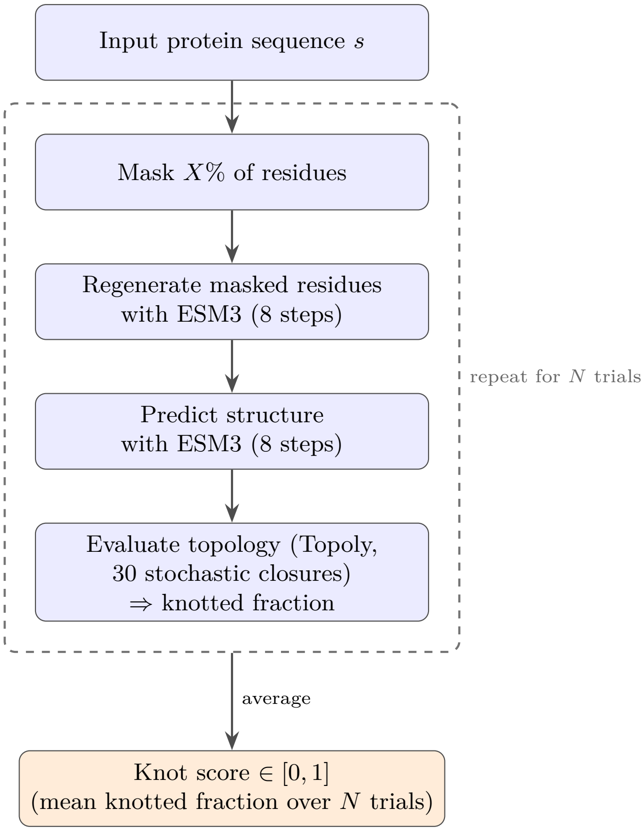


Figure S1: Workflow for the randomized knot score. For a fixed masking level $X$, a single trial masks $X\%$ of the residues, regenerates them with ESM3, predicts the structure of the regenerated sequence, and evaluates its topology with Topoly (30 stochastic closures), yielding the fraction of closures that are knotted. This masking-to-topology pipeline (dashed box) is repeated for $N$ independent trials, and the knotted fractions are averaged to give a continuous knot score in $[0, 1]$. The same per-trial pipeline underlies the random, contiguous, and targeted masking analyses; only the choice of which residues are masked differs.

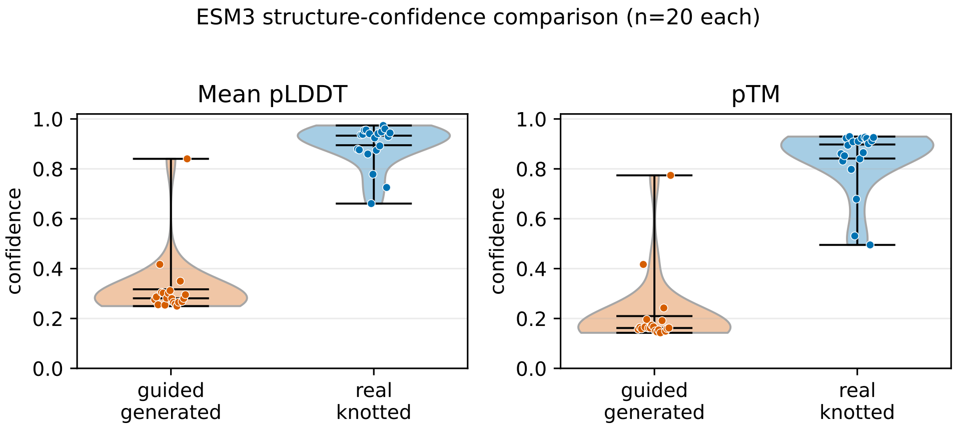


Figure S2: ESM3 structure-confidence comparison for a fresh guided-generation replicate ($n = 20$) and real knotted proteins from the dataset ($n = 20$). Guided generation retained strong topological enrichment but had lower pLDDT and pTM on average than real knotted proteins predicted under the same ESM3-SM structure-prediction pipeline.

Table S1: Targeted masking: core vs. non-core regions ($n = 40$).

| Region masked | % of total seq | Mean knot prob. |
|---|---|---|
| 100% of knot core | ∼47% | 0.659 |
| 100% outside core | ∼53% | 0.828 |
| 100% random (entire seq) | 100% | 0.084 |

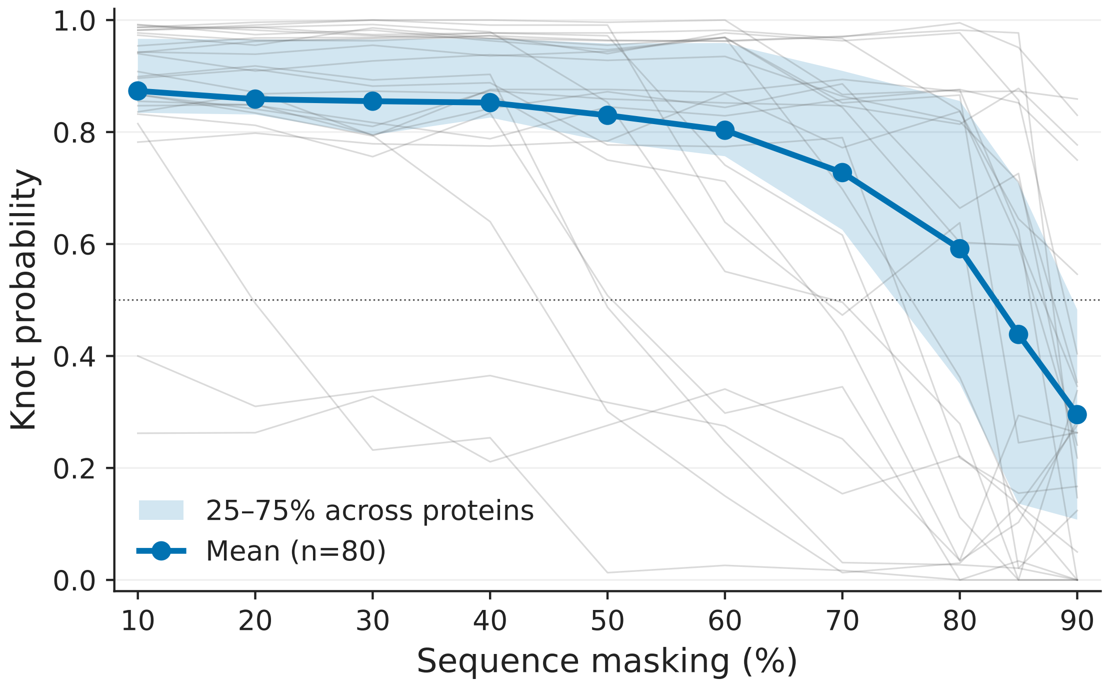


Figure S3: A subset of individual protein transition curves (gray, low opacity) overlaid with the population mean (blue) and the 25–75% inter-quartile range (shaded). The shape of the population curve is dominated by a sharp transition rather than a gradual decline.

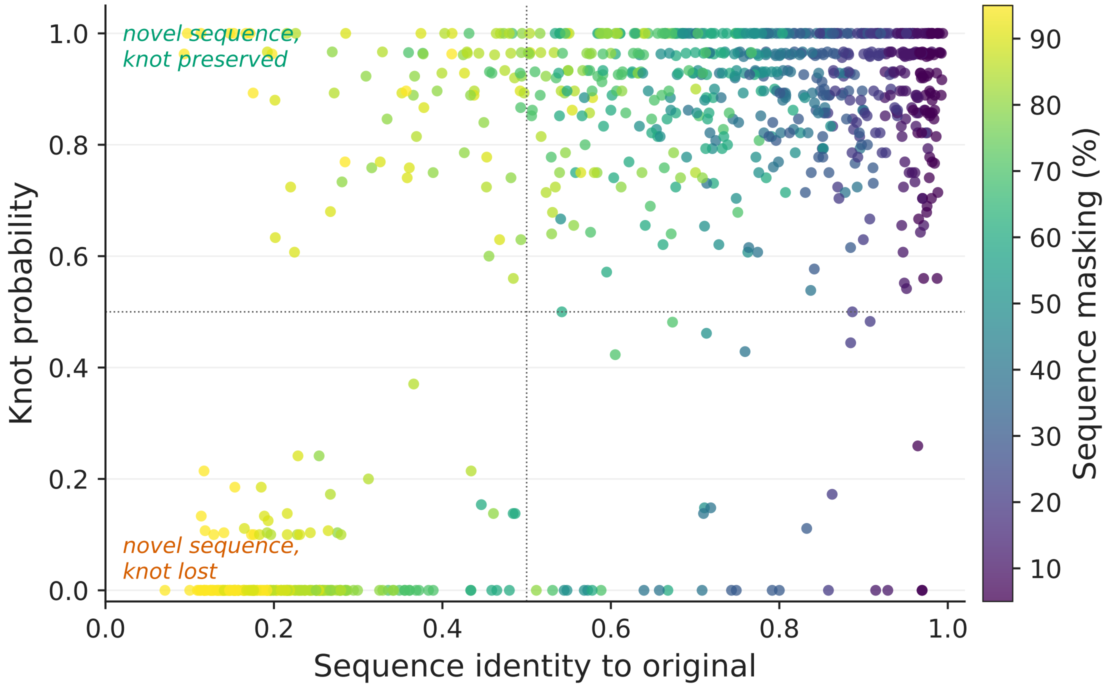


Figure S4: Per-protein sequence identity (after masking and regeneration) versus knot probability. Each point is one (protein, masking level) pair; color encodes the masking percentage.

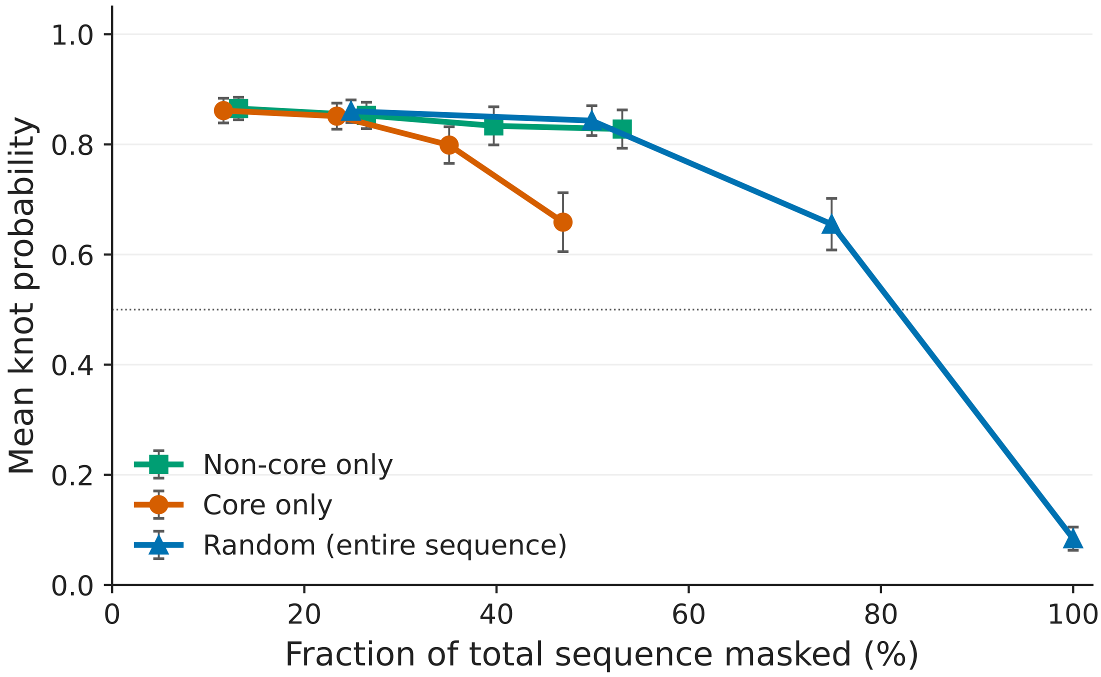


Figure S5: Targeted masking of the knot core (orange) versus non-core regions (green), compared to the random masking baseline (blue). Error bars are standard error of the mean across $n = 40$ proteins. Even at the maximum core-only masking (about 47% of the entire sequence), knot probability stays above 0.65, while random masking of the same fraction drops below 0.35.

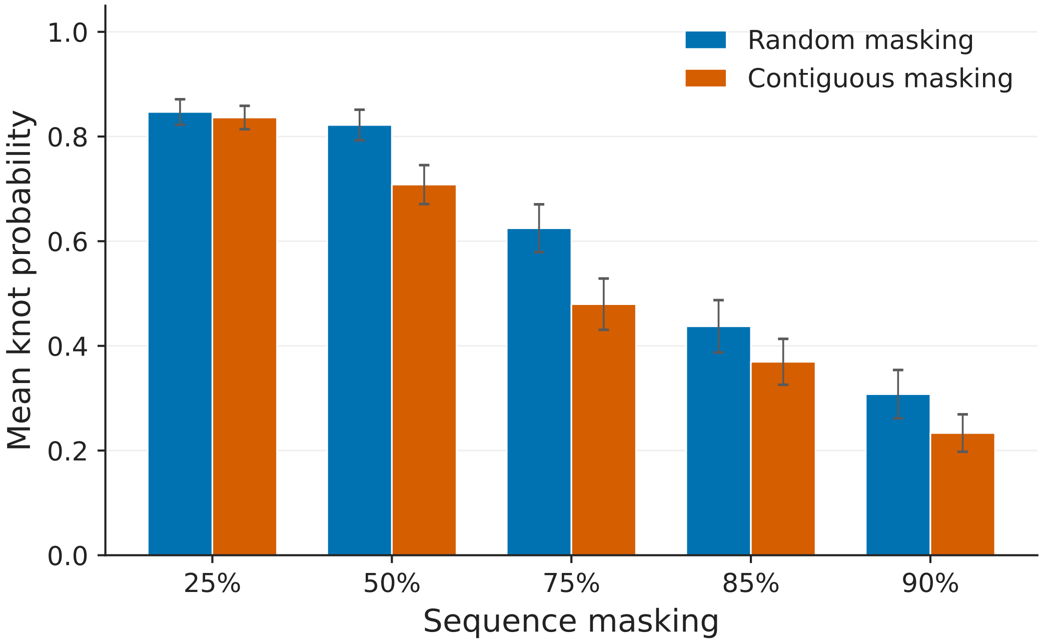


Figure S6: Contiguous masking versus random masking comparison.

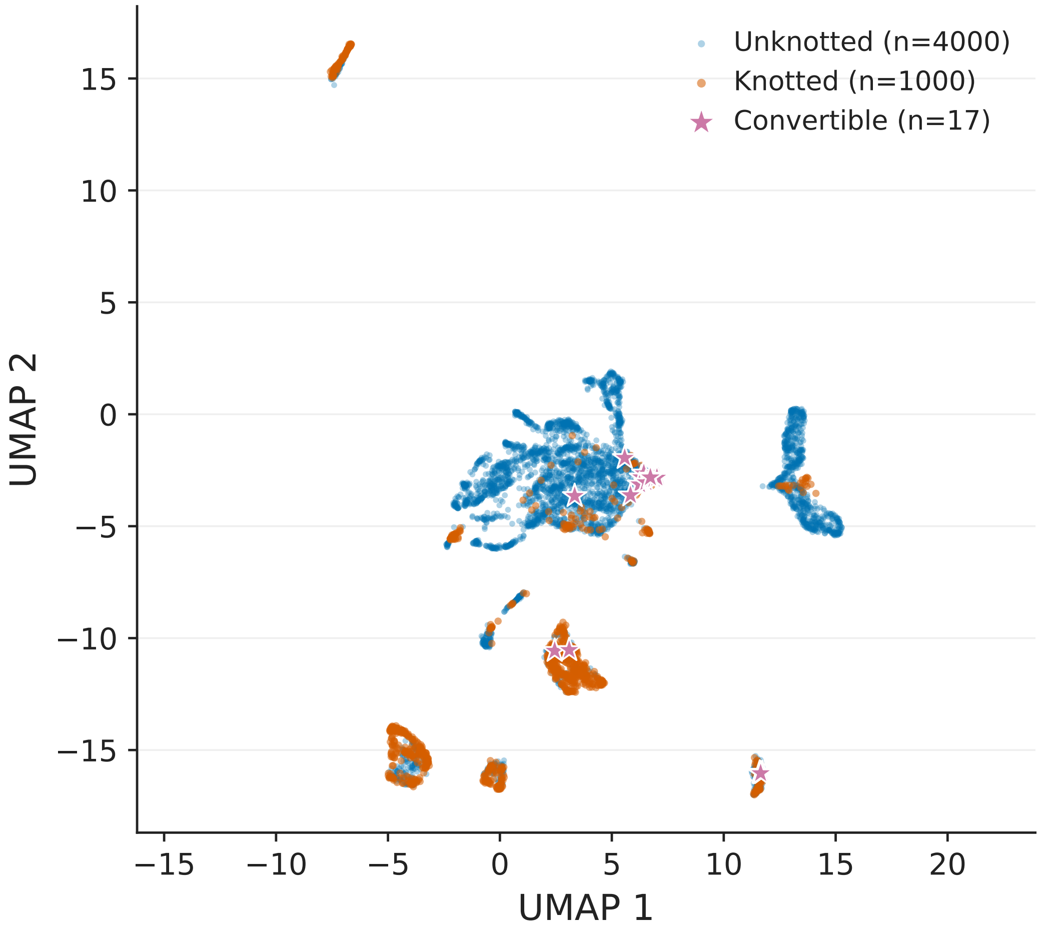


Figure S7: UMAP projection of ESM3 embeddings for 5,000 real proteins. A simple MLP classifier achieves 97.1% accuracy on this representation.

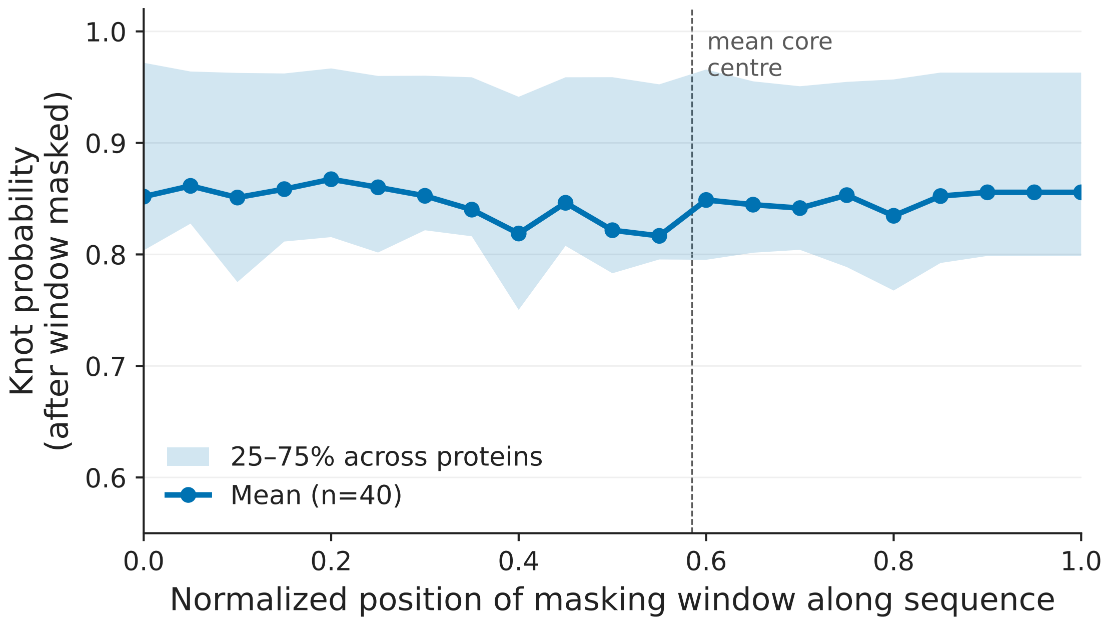


Figure S8: Sliding-window vulnerability profile ($n = 40$, window size 50 residues). The blue line shows the mean knot probability after masking a window centred at each normalized position, with the 25–75% interquartile range shaded. The dashed vertical line marks the mean knot-core centre. The profile is essentially flat, indicating no single position along the sequence is uniquely critical for the knot.

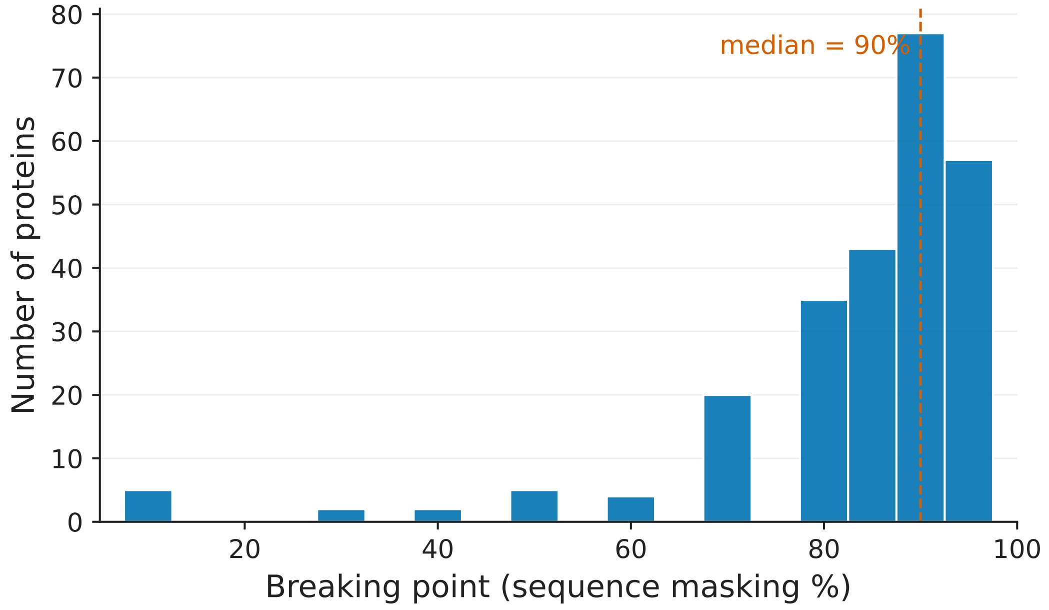


Figure S9: Distribution of breaking points (the smallest masking level at which knot probability drops below 0.5) across 250 knotted proteins. The dashed orange line marks the median (90%), confirming that most proteins retain their knot until very high masking.

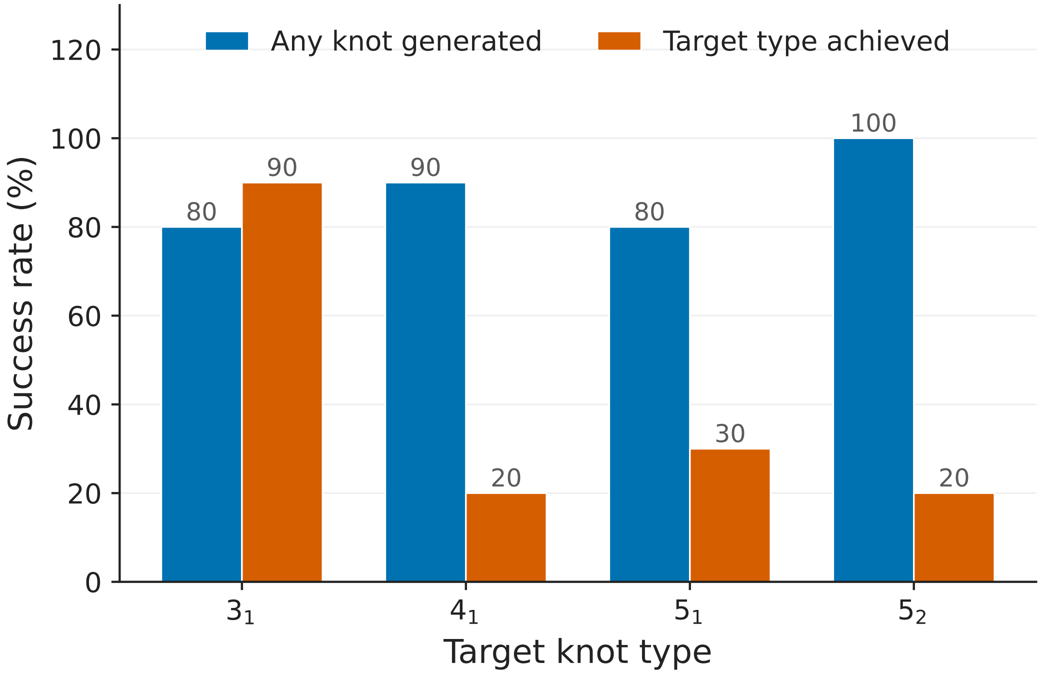


Figure S10: Knot-type-specific guided generation. Blue bars show the rate at which the model generated any knot, orange bars the rate at which it generated the targeted topology specifically. Targeting the trefoil ($3_1$) is highly effective; more complex topologies are produced less reliably and the model often defaults to $3_1$ instead.
